# A T-cell engager platform induces deep plasma cell depletion in autoimmune dry eye

**DOI:** 10.64898/2026.02.06.704311

**Authors:** Jie Ling, Dongliang Wang, Jiangbo Liang, Jingni Li, Qiuling Hu, Qikai Zhang, Xianchai Lin, Zitian Liu, Ting Huang, Ying-Feng Zheng

## Abstract

Bispecific T cell engagers (TCEs) have transformed targeted immunotherapy but remain constrained by *ex vivo* manufacturing and systemic delivery, limiting their application in autoimmune disorders. Here, we introduce the first *in vivo* TCE platform, utilizing a T cell-targeted integration-deficient lentiviral vector (T-IDLE) to reprogram endogenous T cells into engineered T cells secreting BCMA×CD3 TCEs (STCEs) directly *in vivo*. In murine Sjögren’s syndrome-associated dry eye model, a single intravenous administration of T-IDLE_STCE_ achieved deep and sustained depletion of plasmablasts and plasma cells, reversing lacrimal gland inflammation and restoring tear production without detectable off-target transduction. Longitudinal studies in cynomolgus monkeys confirmed translational promise, demonstrating potent plasma cell clearance and favorable safety profiles over 12 weeks post-vector dosing. Mechanistic analyses demonstrated attenuation of Th1/Th17-mediated inflammation and controlled cytokine responses, accompanied by negligible vector integration in non-T cell lineages, thereby substantiating the safety profile of the integration-deficient design. This first-in-class *in vivo* TCE approach represents a manufacturing-free, scalable immunotherapy strategy for autoantibody-mediated diseases.

## Introduction

Sjögren’s syndrome is a chronic autoimmune disease characterized by lymphocytic infiltration of exocrine glands, with xerostomia and keratoconjunctivitis sicca representing its earliest and most frequent clinical manifestations^1-3^. Sjögren’s-syndrome-associated dry eye (SSDE) is a severe, progressive ocular subtype that significantly impairs lacrimal gland function and ocular surface health^3,4^. Current therapeutic approaches are largely palliative^5,6^, and conventional antibody-based therapies lack the capacity to eliminate pathogenic cellular reservoirs in affected tissues or to achieve durable, drug-free remission^7-9^.

Bispecific T cell engagers (TCEs) represent a promising strategy for redirecting endogenous T lymphocytes to selectively recognize and eliminate pathogenic cells^8,10,11^. TCEs are currently under clinical investigation for the treatment of hematologic malignancies and several autoimmune diseases^7,12^. TCEs targeting BCMA×CD3, such as teclistamab, can deplete plasmablasts (PBs) and plasma cells (PCs), the primary source of pathogenic autoantibodies. However, their clinical efficacy varies across autoimmune settings, and serum autoantibody titers in SjS decline only modestly after BCMA×CD3 TCE therapy^7,8^, likely limited by their dependence on the presence and proximity of effector T cells^8,13^.

Genetically arming T cells to constitutively express TCEs may enhance the depth and durability of pathogenic cell depletion^13-15^. Engineered T cells secreting T cell engagers (STCE-T cells) have demonstrated controlled cytokine release upon target engagement^14^, thereby mitigating the excessive cytokine storms commonly associated with CAR-T therapies. This is particularly advantageous for treating inflammatory autoimmune diseases^16-18^.

However, current generation of TCE-engineered T cells relies on complex *ex vivo* manufacturing workflows analogous to CAR-T cell therapy, which are labor-intensive, costly, and associated with systemic and local adverse effects^13,19,20^. Inspired by *in vivo* CAR-T strategies^19^, we developed a novel T cell–targeted, integration-deficient lentiviral vector (T-IDLE) encoding a bispecific antibody that binds CD3 on T cells and bivalently targets BCAM, a surface marker selectively expressed on PBs and PCs^21,22^. BCMA’s absence on naïve and memory B cells, hematopoietic stem cells, and most non-hematopoietic tissues^23^ minimizes off-target depletion of normal immune populations.

In a murine model of SSDE, administration of T-IDLE_STCE_ induced deep and durable depletion of PCs in peripheral blood, lymph nodes, and lacrimal glands, surpassing both conventional TCEs and immunosuppressants in efficacy and durability. Importantly, T-IDLE_STCE_ did not elicit prolonged T cell activation following target clearance, thereby helping to avoid potential early and late toxicities. Likewise, in cynomolgus monkeys, T-IDLE_STCE_ produced durable depletion of PCs, underscoring its translational promise.

In summary, we establish the first-in-class *in vivo* TCE platform optimized for autoimmune disease. It combines the modularity and potency of conventional TCEs while circumventing the limitations of engineered T cell therapy. This approach offers a versatile and clinically scalable strategy for treating autoantibody-mediated autoimmune disorders.

## Results

### Plasma cells are elevated in pSS patients and in the SSDE mouse model

We first analyzed peripheral blood mononuclear cells (PBMCs) from healthy controls (HC) and patients with primary Sjögren’s syndrome (pSS) (Figure 1A). Compared with HC, pSS blood samples showed markedly higher proportions of BCMA⁺ cells among CD45⁺ leukocytes and among CD19⁺ B cells, alongside an increased frequency of circulating PCs (Figure 1B).

**Figure 1.**
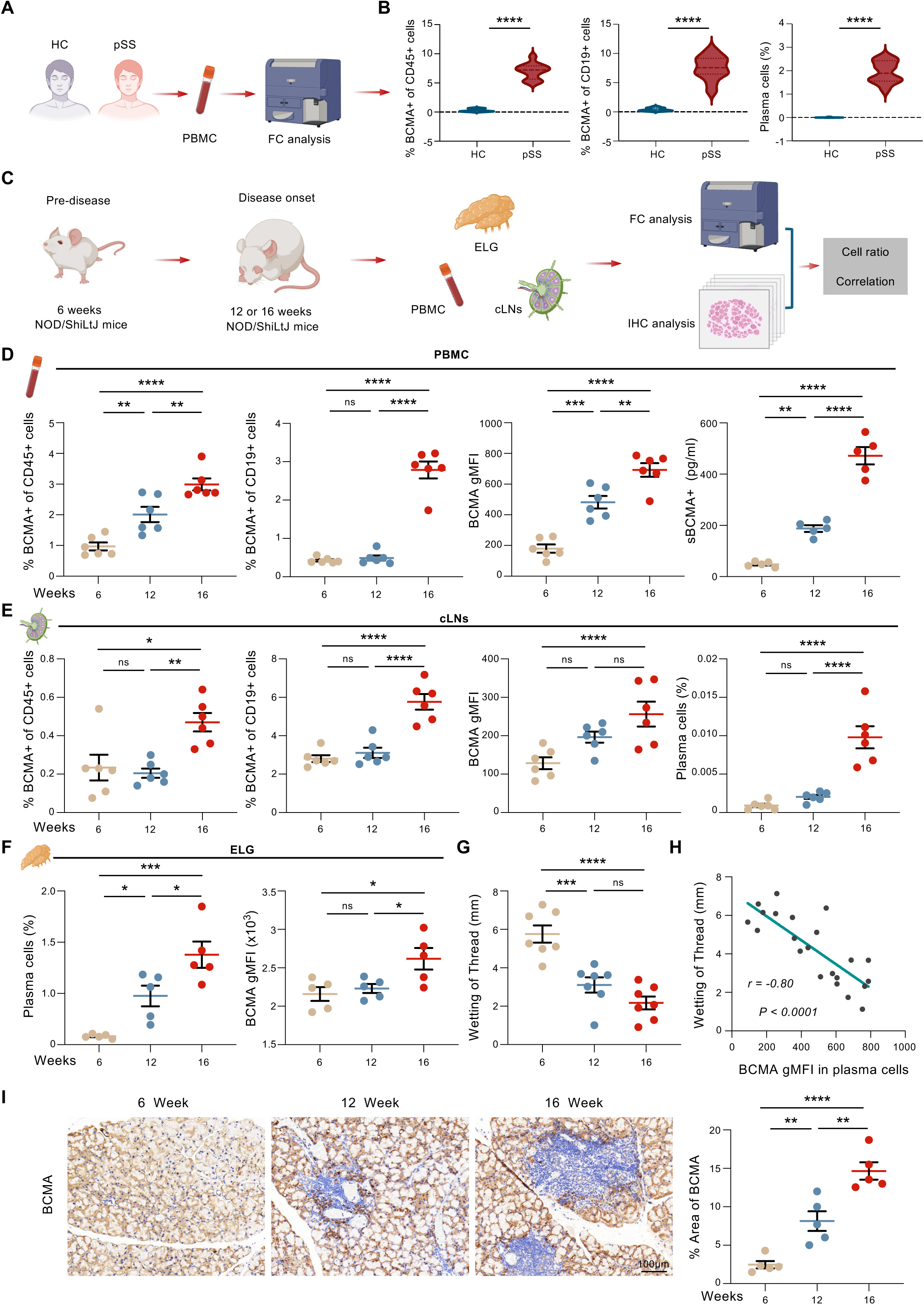
Plasma cells as a therapeutic target in primary SS patients and the mouse SSDE model. (A) Schematic workflow of flow cytometric (FC) analysis of peripheral blood mononuclear cells (PBMCs) from healthy controls (HC) and patients with primary Sjögren’s syndrome (pSS). (B) Frequency of BCMA⁺ cells among CD45⁺ PBMCs (left), BCMA⁺ cells among CD19⁺ B cells (middle), and plasma cells among CD45⁺ PBMCs (right) in HC (n=12) and pSS patients (n=12). (C) Experimental design for analysis of disease progression in the SS mouse model. PBMCs, cervical lymph nodes (cLNs), and extraorbital lacrimal glands (ELGs) were collected at different time points (6, 12, and 16 weeks) by FC and immunohistochemistry (IHC). (D) Flow cytometric quantification of BCMA⁺CD45⁺ and BCMA⁺CD19⁺ cells in PBMCs (left two panels), BCMA gMFI (geometric mean fluorescence intensity) in plasma cells (third panel), and soluble BCMA (sBCMA) levels in serum (right panel) across disease stages (n=5–6 per group). (E) Analysis of cLNs showing frequencies of BCMA⁺CD45⁺ and BCMA⁺CD19⁺ cells, BCMA gMFI in plasma cells, and plasma cell percentage across 6, 12, and 16 weeks (n=6 per group). (F) Plasma cell frequency and BCMA expression in ELGs (n=5 per group). Increased BCMA expression and plasma cell infiltration correlated with disease progression. (G) Phenol red thread test showing tear secretion as a measure of glandular function across disease stages (n=6 per group). (H) Linear regression analysis showing an inverse correlation between BCMA gMFI in plasma cells and tear production (*r* = -0.80, *P* < 0.0001). (I) Representative BCMA IHC staining of ELG sections from mice aged 6-, 12-, and 16-weeks (left), with quantified BCMA⁺ area (right; n=5 per group). The numerical data in (B, D-G, and I) are presented as the mean ± SEM. ns, not significant, *p < 0.05, **p < 0.01, ***p < 0.001, and ****p < 0.0001. A two-tailed unpaired t-test (B) and a one-way ANOVA multiple comparisons test (D-G and I) were used for statistical significance analysis.

To investigate disease progression, we examined a NOD/ShiLtJ mice model of SSDE^24^, collecting PBMCs, cervical lymph nodes (cLNs), and extraorbital lacrimal glands (ELGs) at pre-disease (6 weeks), disease onset (12 weeks), and established disease stages (16 weeks) for flow cytometric and immunohistochemical analysis (Figure 1C). In PBMCs, the frequency of BCMA⁺CD45⁺ cells increased significantly between 6 to 12 weeks and remained elevated at 16 weeks, while BCMA⁺CD19⁺ cells showed no significant change at 12 weeks (Figure 1D). BCMA expression intensity, measured as geometric mean fluorescence intensity (gMFI) on PCs, rose progressively alongside increasing serum sBCMA concentrations (Figure 1D). These dynamic changes likely reflect progressive tissue infiltration of PCs and PBs into lymph nodes and lacrimal glands^2^. Consistently, BCMA expression, a marker of B cell maturation, showed a continuous elevation during disease progression.

Within cervical lymph nodes (cLNs), both BCMA⁺ fractions within CD45⁺ and CD19⁺ compartments and BCMA gMFI on PCs increased from 6 to 16 weeks, with a concomitant rise in PC frequency (Figure 1E). In ELGs, PC infiltration and BCMA expression were significantly elevated at disease onset (12-16 weeks) compared to pre-disease (6 weeks) (Figure 1F). Functional assessment using the phenol red thread test revealed a progressive decline in tear secretion correlating inversely with BCMA gMFI on PBMC PCs (Figures 1G and 1H). Immunohistochemical staining of ELGs confirmed increasing BCMA⁺ cell infiltration over time, as reflected by an growing BCMA⁺ area within affected glands (Figure 1I). These data highlight PC expansion and augmented BCMA expression as characteristic features of SS progression, positioning BCMA as a potential therapeutic target.

### Construction and validation of T cell-targeted integration-deficient lentiviral vectors

Building on the observation that plasma cells accumulate in both SS patients and the SSDE mouse model, where their presence correlates with disease progression, we aimed to develop a targeted delivery system to genetically engineer T cells capable of selectively eliminating these pathogenic cells. To this end, we engineered integration-deficient lentiviral vectors, termed T-IDLE, pseudotyped with an anti-CD3 single-chain variable fragment (scFv) fused to a mutated cocal glycoprotein envelope. This design enables specific binding and transduction of CD3⁺ T cells while avoiding off-target transduction mediated by low-density lipoprotein receptor (LDLR) pathways (Figure 2A).

**Figure 2.**
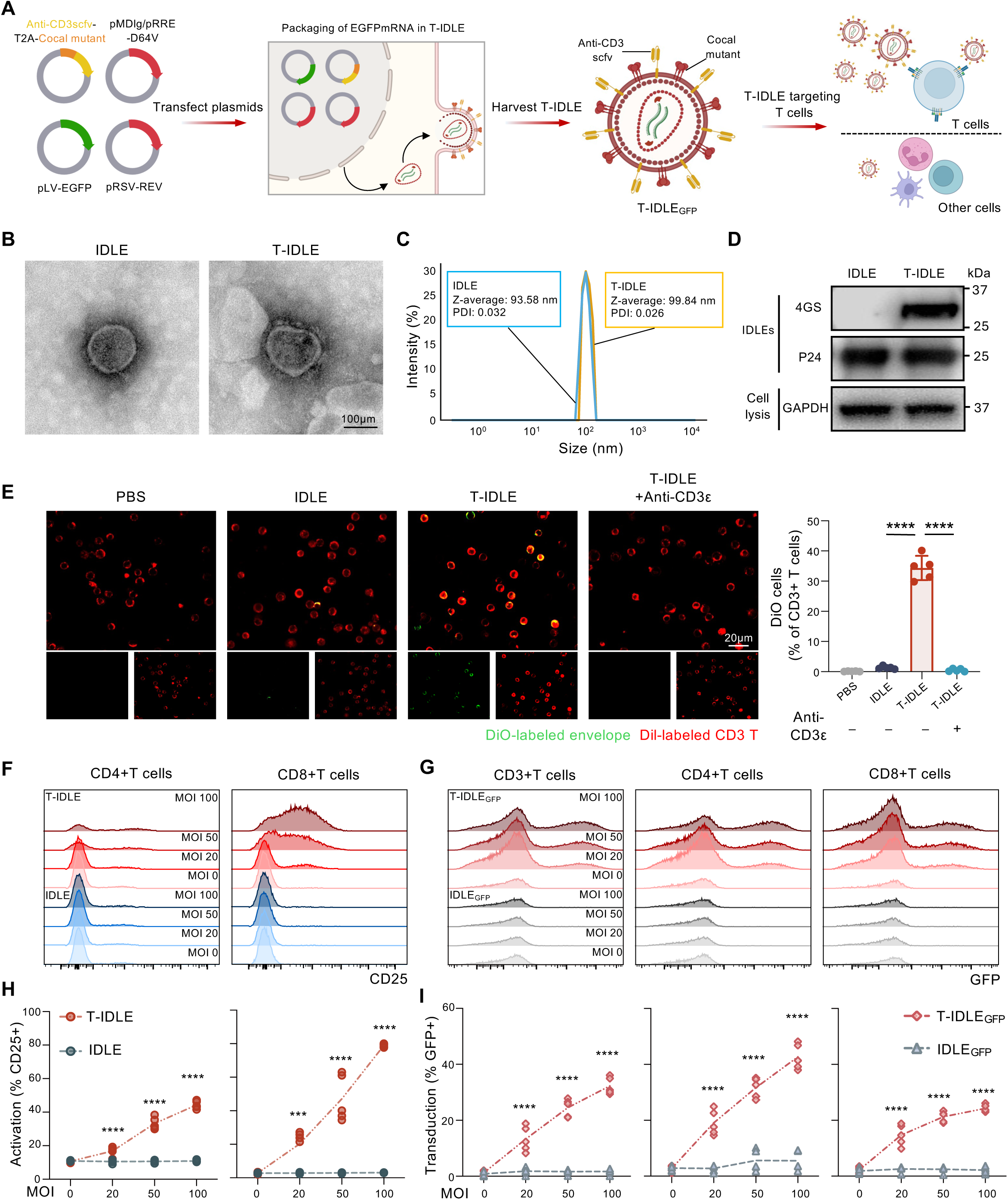
Construction and validation of T cell-targeted integration-deficient lentiviral vectors. (A) Schematic overview of the T-IDLE vector system design. Plasmids encoding anti-CD3 single-chain variable fragment (scFv), cocal glycoprotein mutant, and packaging components were co-transfected to generate T-IDLE particles. The lentiviral envelope displays anti-CD3 scFv to enable specific targeting of CD3⁺ T cells, and includes a mutated cocal glycoprotein to eliminate LDLR binding. (B) Transmission electron microscopy (TEM) images of conventional IDLE and T-IDLE particles. Scale bar, 100 nm. (C) Dynamic light scattering (DLS) measurements of particle size distribution. (D) Western blot detection of lentiviral capsid protein p24 and T-IDLE-associated surface anti-CD3 scFv (anti-G4S linker) in concentrated viral supernatants. (E) Fluorescence microscopy images showing the fusion of DiO-labeled envelopes (green) to CD3⁺ T cells (red) in PBMC samples (n=5 biologically independent samples). Scale bar, 20 nm. (F) FC histograms showing CD25 upregulation (T cell activation marker) in CD4⁺ and CD8⁺ T cells following transduction with T-IDLE or IDLE at increasing MOIs (0, 20, 50, and 100). (G) FC histograms demonstrating GFP expression (as a surrogate of transduction efficiency) in total CD3⁺ T cells, as well as CD4⁺ and CD8⁺ T cell subsets. (H) Quantification of T cell activation (CD25⁺ percentage) induced by T-IDLE and IDLE at varying MOIs (n=5 biologically independent samples). (I) Quantification of GFP⁺ transduction efficiency in CD3⁺, CD4⁺, and CD8⁺ T cells (n=5 biologically independent samples). The numerical data in (E, H, and I) are presented as the mean ± SEM. ****p < 0.0001. A one-way ANOVA multiple comparisons test (E) and a two-way ANOVA with multiple comparisons test (H and I) were used for statistical significance analysis.

Transmission electron microscopy (TME) revealed both conventional IDLE and T-IDLE particles exhibited comparable spherical morphology without discernible ultrastructural differences (Figure 2B). Dynamic light scattering (DLS) analysis confirmed a modest increase in the hydrodynamic diameter of T-IDLE particles, potentially attributable to the display of anti-CD3 scFv on their surface (Figure 2C). Western blotting (WB) of concentrated vector supernatants verified equivalent levels of p24 capsid protein in IDLE and T-IDLE preparations, with specific detection of anti-CD3 scFv on T-IDLE envelopes (Figure 2D).

Fluorescence microscopy revealed efficient fusion of green-labeled T-IDLE envelopes with red-labeled CD3⁺ T cells. Pretreatment of target cells with Anti-CD3ε antibodies significantly reduced fusion efficiency, confirming CD3-dependent targeting specificity (Figure 2E). Flow cytometric analyses revealed that T-IDLE transduction induced dose-dependent upregulation of the activation marker CD25 on both CD4⁺ and CD8⁺ T cells across multiplicities of infection (MOIs) ranging from 0-100, significantly outperforming non-targeted IDLE vectors (Figure 2F). Similarly, GFP reporter expression, as a surrogate marker for transduction efficiency, was markedly higher in total CD3⁺, CD4⁺, and CD8⁺ T cell populations transduced with T-IDLE compared to IDLE at all tested MOIs (Figure 2G). Quantitatively, T-IDLE elicited up to 80% CD25⁺ activation in CD8⁺ T cells at an MOI 100, with a similar activation profile observed in CD4⁺ T cells (Figure 2H). Overall transduction efficiency reached ∼36.1% GFP⁺ cells within CD3⁺ populations, with CD4⁺ T cells displaying higher uptake compared to CD8⁺ counterparts (Figure 2I). These findings validate T-IDLE as a stable, T cell-specific vector platform capable of efficient gene delivery coupled with controlled T cell activation kinetics.

### Generation and functional characterization of T-IDLE-induced STCE-T cells

Having validated the specificity of the T-IDLE vector for T cell transduction, we employed T-IDLE vectors carrying TCE cargo to engineer T cells capable of secreting bispecific engagers (STCE-T cells) for targeted depletion of PCs and pathogenic autoantibodies (Figure 3A). Co-culture of STCE-T cells generated via T-IDLE with BCMA-expressing target cells revealed consistent secretion of functional TCEs (Figure 3B). Flow cytometric analysis confirmed rubost TCE-mediated conjugate formation, with approximately 60% of cells in T-IDLE_STCE_-treated samples exhibiting CD3⁺BCMA⁺ double positivity, a significant increase compared to PBS, IDLE, or T-IDLE controls (Figure 3C). Confocal microscopy of co-cultures further illustrated dynamic TCE-mediated interactions, with green-labeled CD3⁺ STCE-T cells forming intimate contacts with red-labeled A20^BCMA^ target cells (Figure 3D).

**Figure 3.**
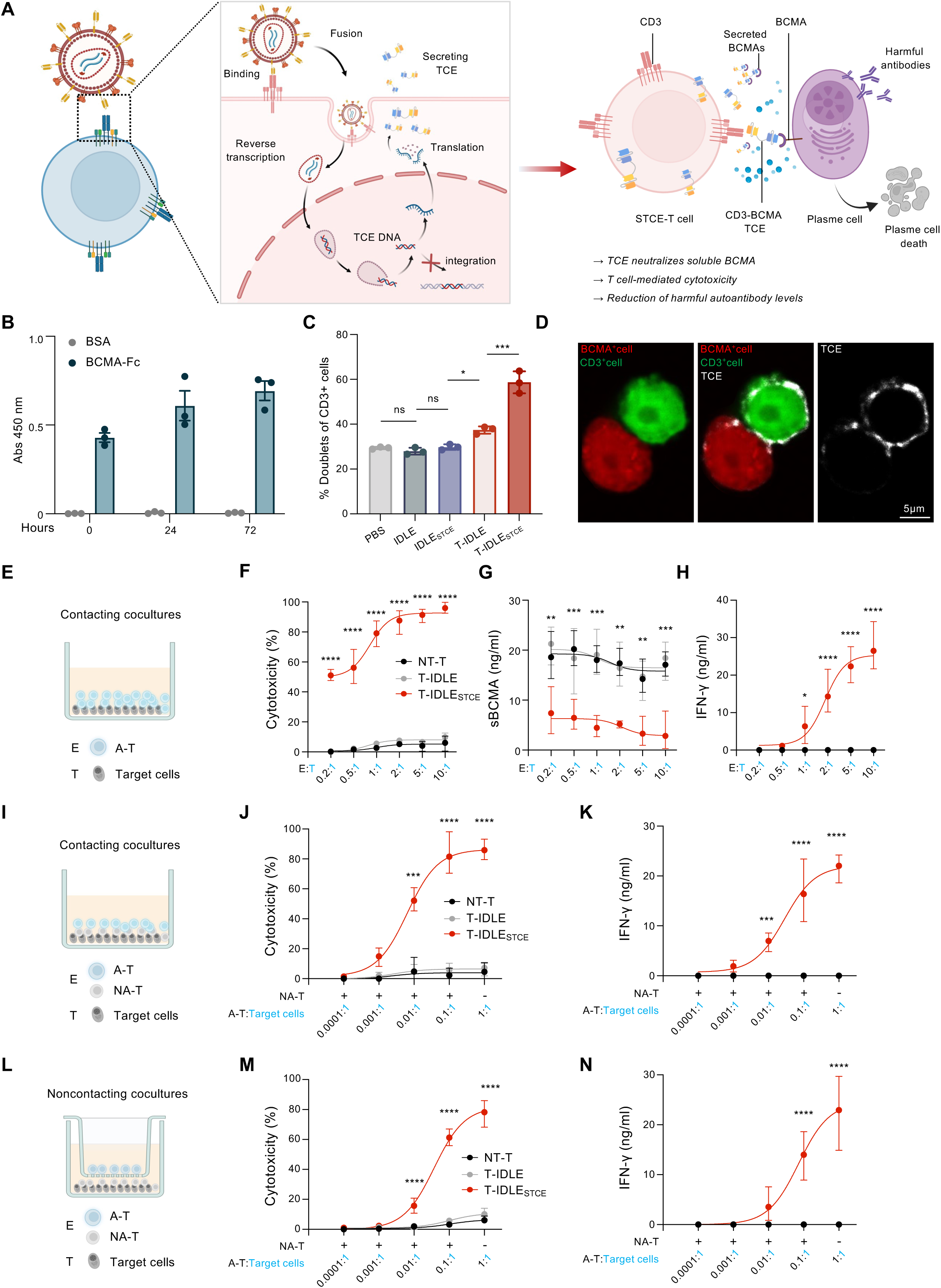
Generation and functional characterization of T-IDLE induced STCE-T cells. (A) Schematic of STCE-T cell generation following T-IDLE transduction. T-IDLE binds and fuses with CD3⁺ T cells, delivering TCE-encoding sequences via reverse transcription without genomic integration. Transduced STCE-T cells secrete TCEs, which neutralize soluble BCMA (sBCMA) and mediate plasma cell killing through TCE formation. (B) Quantification of TCE binding activity by ELISA. Soluble TCE secretion from STCE-T cells generated via T-IDLE transduction was assessed by ELISA at 0, 24, and 72 hours after co-culture with BCMA-expressing target cells. Binding to recombinant BCMA-Fc was measured, with BSA serving as a negative control (n=3 biologically independent samples). (C) FC analysis of CD3⁺BCMA⁺ double-positive cells after transduction with indicated components (n=3 biologically independent samples). (D) Confocal microscopy images of STCE-T cells co-cultured with BCMA⁺ target cells (CD3, green; BCMA, red; TCE, white). Scale bar, 5 µm. (E) Schematic of direct contact co-culture assay. Effector T cells (E) and BCMA⁺ target cells (T) were co-incubated at defined E:T ratios. (F-H) Functional assays showing dose-dependent cytotoxicity (F), reduction in sBCMA levels (G), and IFN-γ secretion (H) in STCE-T cells compared with NT-T (nontransduced T cells) and T-IDLE controls across E:T ratios (n=3 biologically independent samples). (I) Schematic of direct contact co-culture assay in which engineered activated T cells are mixed with non-activated T cells (NA-T) and co-cultured with BCMA⁺ targets at the indicated A-T:target ratios. (J and K) In direct contact co-culture assays, cytotoxicity (J) and IFN-γ (K) increased as the fraction of A-T rises, indicating that secreted STCE armed bystander T cells (n=3 biologically independent samples). (L) Schematic of non-contacting transwell assay isolating A-T from targets to test for diffusion-mediated activity of secreted TCE. (M and N) In non-contacting assays, cytotoxicity (M) and IFN-γ (N) scaled with the A-T:target ratio for T-IDLE_STCE_, demonstrating that soluble STCE is sufficient to drive killing and cytokine production without cell-cell contact (n=3 biologically independent samples). The numerical data in (B, C, F-H, J, K, M, and N) are presented as the mean ± SEM. ns, not significant, *p < 0.05, **p < 0.01, ***p < 0.001, and ****p < 0.0001. A one-way ANOVA multiple comparisons test (C) and a two-way ANOVA comparisons test (F-H, J, K, M, and N) were used for statistical significance analysis.

In direct-contact cytotoxicity assays (Figure 3E), STCE-T cells exhibited potent, dose-dependent lysis of BCMA⁺ targets, reaching up to ∼90% target cell killing at an effector-to-target (E:T) ratio of 10:1 (Figure 3F). This cytotoxicity coincided with significant reduction in sBCMA levels (∼80% decrease) (Figure 3G) and elevated IFN-γ secretion, reaching up to approximately 35 pg/mL at the highest tested E:T ration (10:1) (Figure 3H).

To evaluate the potential for bystander T cell activation via soluble TCE, mixed co-cultures of activated T cells (A-T) and non-activated T cells (NA-T) were established at varying A-T:target ratios (Figures 3I-3K). Both cytotoxicity and IFN-γ secretion correlated positively with the proportion of activated effector T cells, reaching ∼80% target lysis and ∼20 pg/mL IFN-γ at a 0.1:1 A-T:NA-T ratio (Figures 3J and 3K), demonstrating the capacity of secreted TCEs to arm bystander T cells effectively. In non-contacting transwell assays separating A-T from targets to assess diffusible TCE activity (Figure 3L), STCE-T cells retained significant cytotoxic and IFN-γ release in a ratio-dependent manner, confirming that diffusible TCE molecules suffice to redirect T cell cytotoxicity without direct cell-cell contact (Figures 3M and 3N). These data establish T-IDLE induced STCE-T cells as highly functional, BCMA-specific effectors that effectively neutralize sBCMA, mediate PC lysis, and recruit bystander T cells via secreted TCE, supporting their therapeutic potential in SS.

### *In vivo* t-cell targeting specificity of T-IDLEs

To assess *in vivo* specificity and transduction efficiency, we administered T-IDLE_GFP_ (Figure 4A) or T-IDLE_STCEGFP_ (encoding the anti-BCMA × anti-CD3 TCE linked via a 2A linker to GFP) (Figure 4B) intravenously to wild-type mice and monitored peripheral blood transduction over 21 days. Flow cytometric analysis revealed GFP expression exclusively in CD3⁺ T cells following T-IDLE_GFP_ treatment, with GFP⁺ populations emerging by day 7 and peaking at day 14, whereas PBS, non-targeted IDLE, and IDLE_GFP_ control groups showed negligible signals (Figure 4C). Quantification confirmed dose-dependent transduction efficiencies, reaching ∼60% GFP⁺ CD3⁺ T cells at day 14 in T-IDLE_GFP_ treated groups, which returned to baseline by day 21 (Figure 4D). Off-target transduction was minimal, as reflected by negligible GFP expression in CD3^-^ cell populations (Figure 4E).

**Figure 4.**
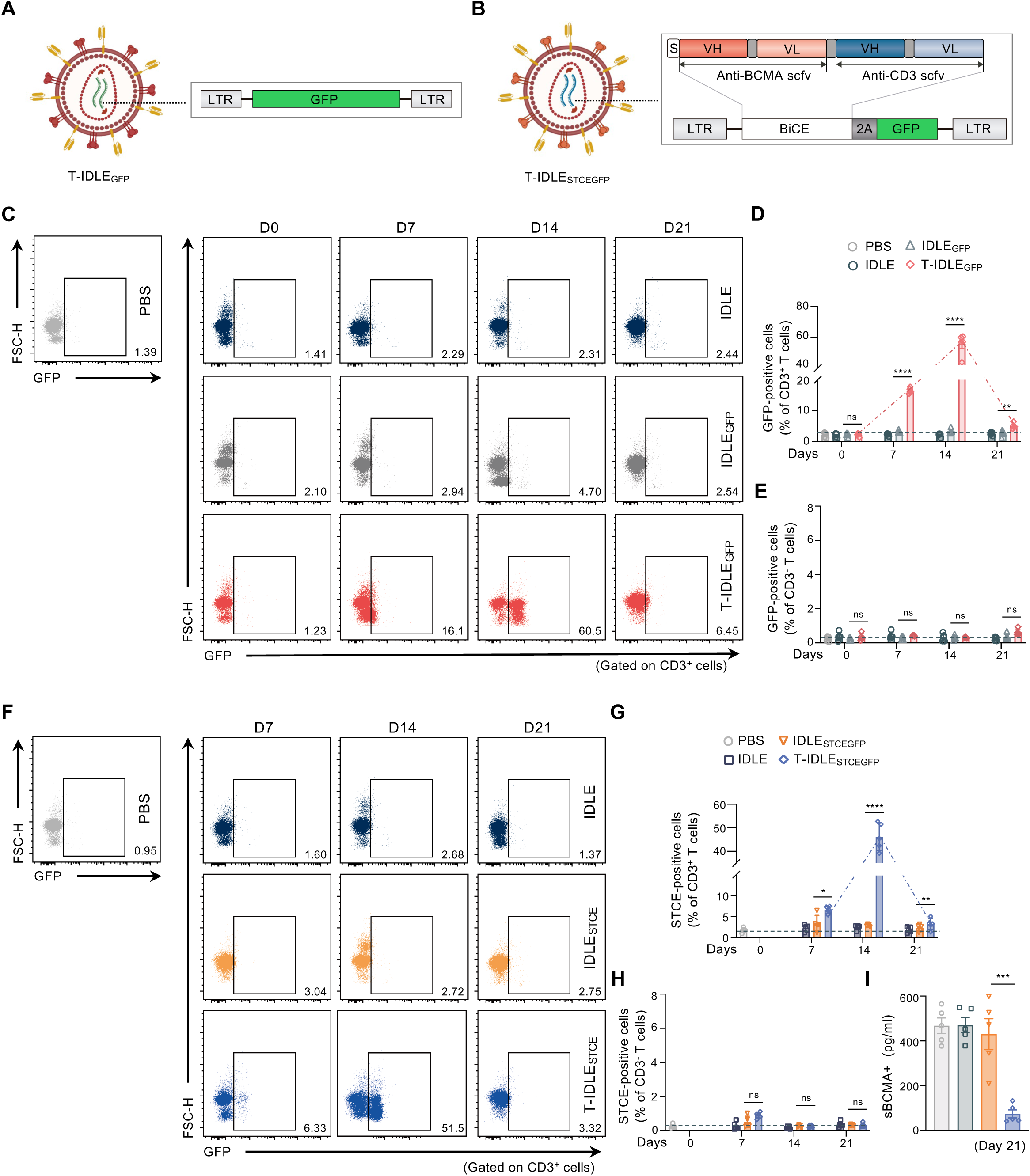
*In vivo* T-cell targeting specificity of T-IDLEs. (A-B) Schematic diagrams of T-IDLE vectors used for *in vivo* studies. T-IDLE_GFP_ (A) encodes GFP alone to assess T cell transduction efficiency; T-IDLE_STCEGFP_ (B) encodes a TCE construct (anti-BCMA × anti-CD3 scFv) and GFP via a 2A linker to generate functional STCE-T cells *in vivo*. (C) Representative FC plots showing GFP⁺ CD3⁺ T cells in PBMCs from mice treated with PBS, IDLE, IDLE_GFP_, or T-IDLE_GFP_ over time (Day 0, 7, 14, 21). (D) Quantification of GFP⁺ CD3⁺ T cells in peripheral blood over time (n=5 per group). (E) Quantification of GFP⁺ cells among off-target populations (n=5 per group). (F) Representative FC plots showing STCE-T cells (GFP⁺CD3⁺) in PBMCs after injection of PBS, IDLE, IDLE_STCEGFP_, or T-IDLE_STCEGFP_ over time (Day 0, 7, 14, 21). (G) Quantification of STCE-T cells over time (n=5 per group). (H) Quantification of STCE-T cells among off-target populations (n=5 per group). (I) Measurement of serum sBCMA levels at day 21 post-injection (n=5 biologically independent samples). The numerical data in (D, E, and G-I) are presented as the mean ± SEM. ns, not significant, *p < 0.05, **p < 0.01, ***p < 0.001, and ****p < 0.0001. A one-way ANOVA multiple comparisons test (D, E, and G-I) was used for statistical significance analysis.

Similarly, T-IDLE_STCEGFP_ administration induced GFP⁺ CD3⁺ populations in PBMCs, begining at day 7 with maximal transduction at day 14, absent in PBS, IDLE, or IDLE_STCEGFP_ control groups (Figure 4F). Transduction efficiency peaked at ∼50% STCE-T cells by day 14, with transient kinetics mirroring those observed with T-IDLE_GFP_ (Figure 4G). Off-target transduction remained low (Figure 4H). Notably, serum sBCMA levels were significantly reduced at day 21 in T-IDLE_STCEGFP_-treated mice (∼50 pg/mL versus ∼500 pg/mL in controls), consistent with TCE-mediated BCMA neutralization (Figure 4I). These results demonstrate that T-IDLE vectors enable highly specific and transient programming of endogenous T cells *in vivo*, with minimal off-target effects.

### T-IDLE_STCE_ treatment induces deep plasma cell depletion in the SSDE model

We next evaluated the capacity of *in vivo* T-IDLE_STCE_ engineering to access disease-relevant tissues and deplete PCs in the SSDE model. A single intravenous dose of T-IDLE_STCEGFP/STCELuc_ or control treatments (PBS, IDLE, or IDLE_STCEGFP/STCELuc_) was administered to 10-week-old SSDE mice at disease onset, followed by longitudinal assessment of STCE-T cell infiltration and PC depletion in PBMCs, ELGs, and cLNs (Figure 5A).

**Figure 5.**
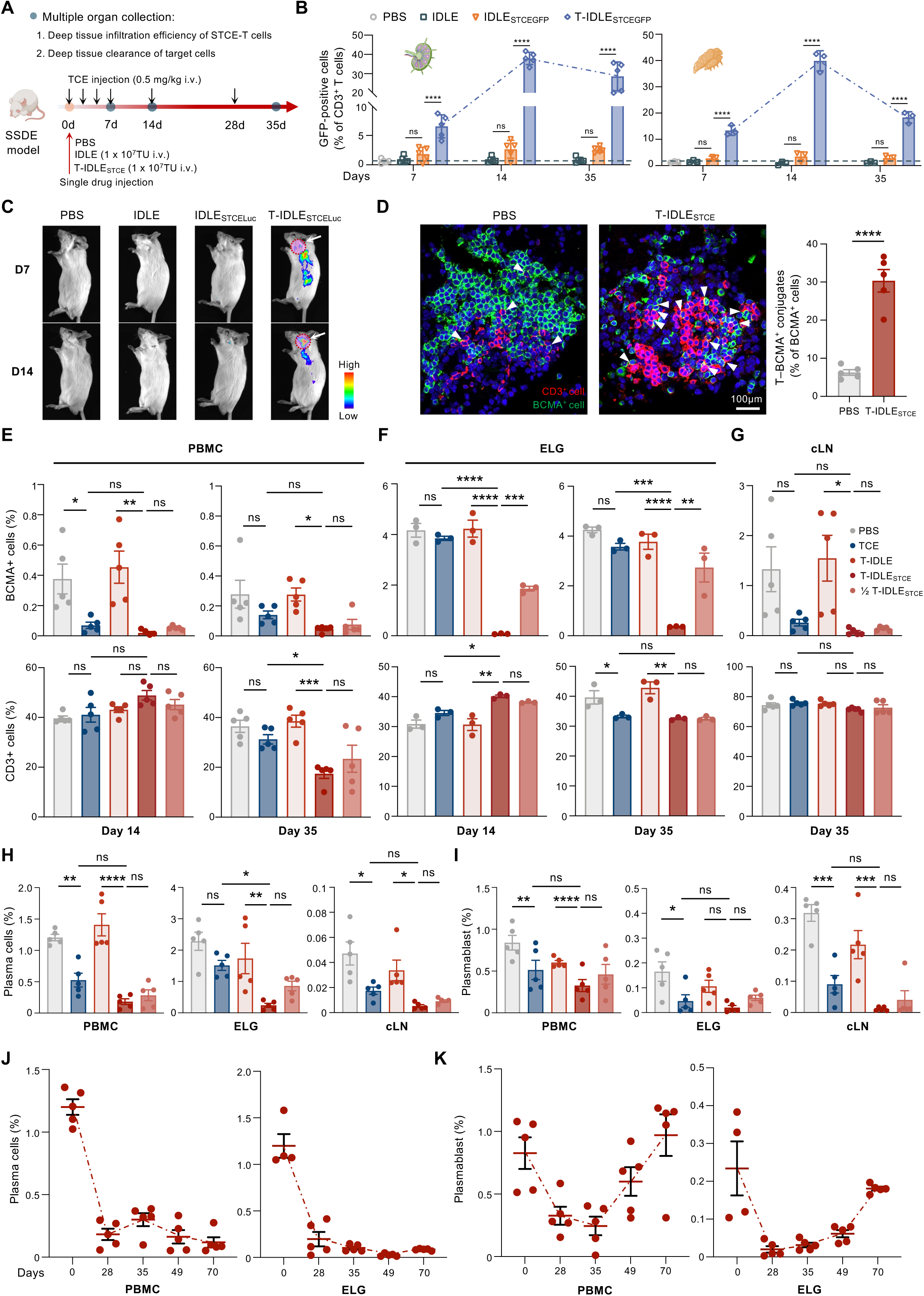
T-IDLE_STCE_ treatment induces deep plasma cell depletion in SSDE model. (A) Schematic of experimental timeline. T-IDLE_STCE_ (1 × 10^7^ TU) or other treatments were administered to 10-week-old SSDE mice. Peripheral blood, cLNs, and ELGs were collected to assess STCE-T cell infiltration and PC clearance. (B) Quantification of GFP⁺ CD3⁺ T cells in ELGs (left) and cLNs (right) over time (Day 7, 14, 35; n=5 or 3 per group). (C) Representative *in vivo* bioluminescence images at day 7 and 14 showing tissue distribution of STCE-T cells. (D) Representative immunofluorescence images of ELGs stained for BCMA (green), CD3 (red), and nuclei (DAPI, blue). Right, quantification of T–BCMA^+^ conjugates among all BCMA^+^ cells (n=5 per group). Scale bar, 100 µm. (E-G) FC analysis of BCMA⁺ and CD3⁺ cell frequencies in PBMC (E), ELGs (F), and cLNs (G) at day 14 or day 35 post-treatment (n=5 or 3 per group). (H) Quantification of plasma cells in PBMCs, ELGs, and cLNs at day 35 (n=5 or 3 per group). (I) Quantification of plasmablasts in PBMCs, ELGs, and cLNs at day 35 (n=5 or 3 per group). (J-K) Longitudinal tracking of plasma cells (J) and plasmablasts (K) in PBMCs and ELGs up to 70 days post-treatment. The numerical data in (B and D-K) are presented as the mean ± SEM. ns, not significant, *p < 0.05, **p < 0.01, ***p < 0.001, and ****p < 0.0001. A two-tailed unpaired t-test (D) and a one-way ANOVA multiple comparisons test (B and E-I) were used for statistical significance analysis.

Flow cytometry revealed that GFP⁺ CD3⁺ STCE-T cells infiltrated ELGs and cLNs, peaking at approximately 40% of CD3⁺ T cells in ELGs by day 14, with a similar infiltration pattern in cLNs (Figure 5B). *In vivo* bioluminescence imaging confirmed STCE-T cells distribution and luciferase signals localized to ELG regions at day 7, which persisted focally at day 14 in T-IDLE_STCELuc_-treated mice but were minimal in controls (Figure 5C).

Immunofluorescence microscopy of ELGs at day 7 showed extensive colocalization of CD3⁺ cells (red) with BCMA⁺ cells (green), forming immune conjugates in T-IDLE_STCE_-treated mice (Figure 5D). Quantification indicated ∼30% of BCMA⁺ cells engaged in T-BCMA⁺ conjugates in T-IDLE_STCE_ groups (Figure 5D). Flow cytometric analysis at day 14 revealed profound reductions in BCMA⁺ cells across compartments (PBMCs and ELGs) in T-IDLE_STCE_-treated mice, with concurrent increases in CD3⁺ T cell frequencies (Figures 5E and 5F). By day 35, PC depletion persisted, with BCMA⁺ cell frequencies remaining low in PBMCs (∼0.1%), ELGs (∼0.3%), and cLNs (∼0.2%), alongside reduced CD3⁺ T cell proportions in both full-and half-dose T-IDLE_STCE_ groups (Figures 5E-5G). The reduction in CD3⁺ T cell proportions may be attributable to decreased autoantibody secretion and higher TCE concentrations masking CD3 epitopes on T cells, mitigating potential side effects arising from excessive T cell proliferation and activation^14^.

Detailed analysis at day 35 confirmed sustained depletion of both PCs and PBs across PBMCs, ELGs, and cLNs following T-IDLE_STCE_ treatment (Figures 5H and 5I). Longitudinal monitoring demonstrated progressive PC clearance in PBMCs and ELGs, reaching near-complete depletion by day 35 and maintaining low levels (<0.5%) through day 70 (Figure 5J), with PB frequencies following a similar trajectory (Figure 5K). These findings underscore the capacity of a single intravenous T-IDLE_STCE_ dose to mediate profound and durable depletion of pathogenic plasma cells in disease-relevant tissues. Our observation aligns with an immune reset that facilitates restoration of immune homeostasis.

### T-IDLE_STCE_ treatment reduces inflammation by modulating T-cell function

Current TCEs have shown suboptimal performance in various malignancies and autoimmune diseases^7,8^, largely due to their rapid immune-mediated clearance from peripheral blood following direct injection or premature exhaustion of effector T cells and PCs within lymph nodes^9,25-28^. This limits their effective penetration into affected tissues. In contrast, our T-IDLE treatment enables in-situ engineering of endogenous T cells into continuous factories producing TCEs, facilitating targeted delivery to affected sites via infiltration of STCE-T cells.

To evaluate the therapeutic impact of T-IDLE_STCE_ in autoimmune microenvironments, we employed the SSDE model to study immune modulation in lymph nodes, lacrimal glands, and corneal tissues. Flow cytometric analysis of CD4⁺ T cell subsets in cLNs at day 14 and 35 (Figure 6A) post-treatment revealed that T-IDLE_STCE_ markedly reduced the frequency of IFN-γ⁺ CD4⁺ Th1 cells compared to SSDE controls, decreasing from ∼2% to ∼0.2% at day 14 and sustaining low levels (∼0.5%) at day 35. In contrast, TCE injection and convention eye drop treatments yielded partial or no effects (Figure 6B). Similarly, IL17A⁺ CD4⁺ Th17 cells were significantly diminished in T-IDLE_STCE_ groups, from ∼1.4% in SSDE to ∼0.2% following T-IDLE_STCE_ treatment at both day 14 and 35, with minimal changes in controls (Figure 6C).

**Figure 6.**
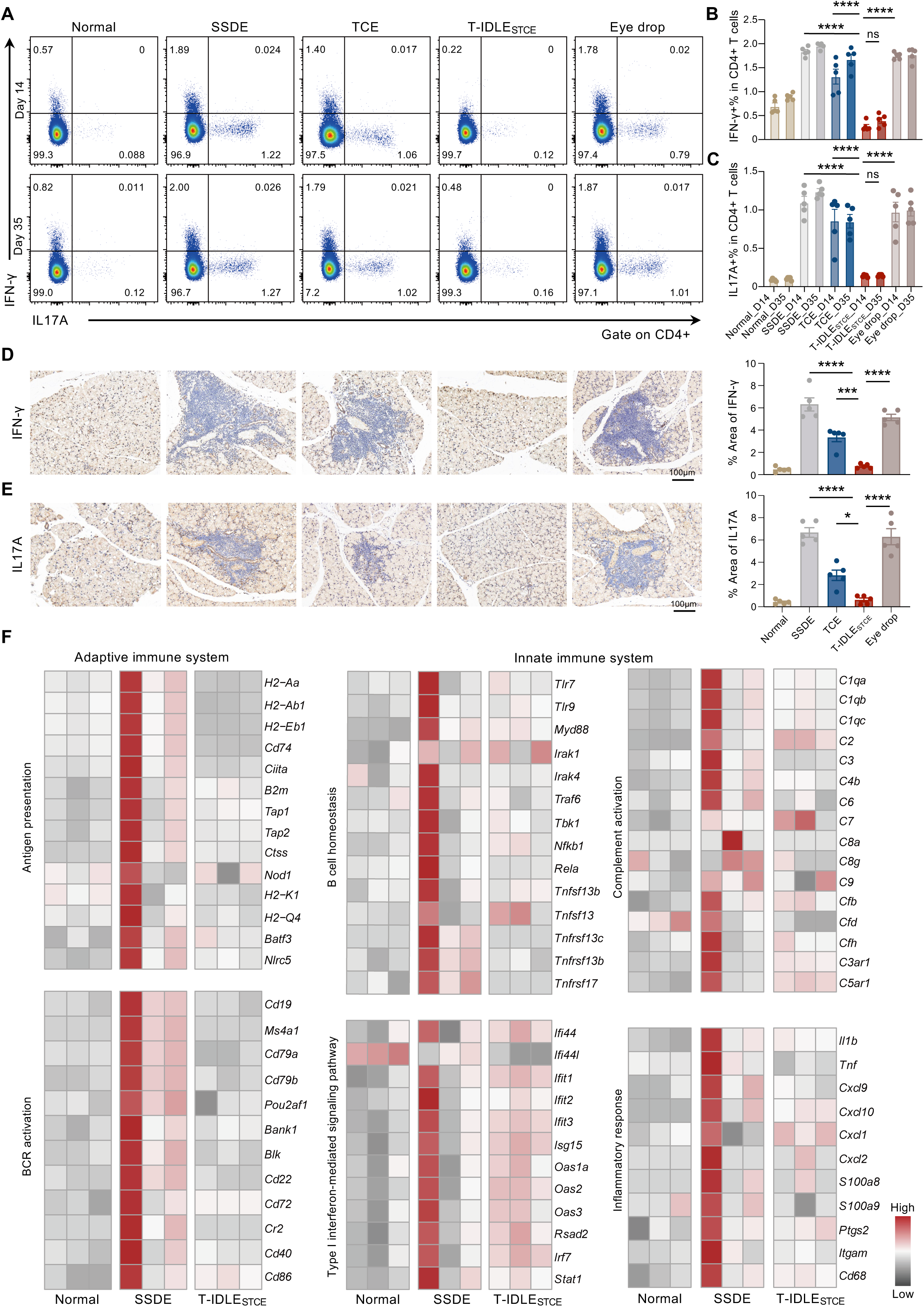
T-IDLE_STCE_ treatment reduces inflammation by modulating T-cell function. (A) Representative FC plots showing frequencies of IFN-γ⁺ and IL17A⁺ CD4⁺ T cells in lacrimal glands at day 14 and 35 across treatment groups. (B) Quantification of IFN-γ⁺ CD4⁺ T cells at day 14 and 35 (n=5 per group). (C) Quantification of IL17A⁺ CD4⁺ T cells at day 14 and 35 (n=5 per group). (D) Representative IHC images for IFN-γ in ELGs and quantification of IFN-γ-positive area (% of tissue) at day 35 (n=5 per group). Scale bar, 100 µm. (E) Representative IHC images for IL17A in ELGs and quantification of IL17A-positive area (% of tissue) at day 35 (n=5 per group). Scale bar, 100 µm. (F) Transcriptomic profiling heatmap of gene expressions associated with antigen presentation, BCR activation, B cell homeostasis, type I interferon-mediated signaling pathway, complement activation, and inflammatory response in ELGs across treatment groups (n=3). The numerical data in (B-E) are presented as the mean ± SEM. ns, not significant, *p < 0.05, ***p < 0.001, and ****p < 0.0001. A one-way ANOVA multiple comparisons test (B-E) was used for statistical significance analysis.

Immunohistochemical analyses of ELGs at day 35 corroborated these findings, revealing sparse IFN-γ staining in T-IDLE_STCE_-treated mice relative to dense infiltrates in control groups, with positive staining areas reduced from ∼6% to ∼1% (Figure 6D). IL17A expression followed a parallel pattern, dropping from ∼8% in SSDE to ∼1% following treatment (Figure 6E).

Transcriptomic profiling of ELGs further elucidated the immunomodulatory effects, with heatmaps showing downregulation of genes involved in antigen presentation (e.g., *H2-Aa*, *Cd74*), B cell signaling and homeostasis (e.g., *Cd19*, *Batf3*), type I interferon pathways (e.g., *Ifit1*, *Isg15*), complement activation (e.g., *C1qa*, *C3*), and inflammatory responses (e.g., *Il1b*, *Cxcl9*) in T-IDLE_STCE_-treated mice compared with SSDE controls. These expression patterns approached those observed in normal controls, indicating restoration of immune homeostasis (Figure 6F). These data indicate that T-IDLE_STCE_-mediated PC ablation attenuates Th1/Th17-driven inflammation and restores immune homeostasis in lymphoid and glandular tissues.

SSDE progression disrupted corneal homeostasis, characterized by elevated reactive oxygen species (ROS), increased expression of pro-inflammatory and epithelial stress markers including interleukin-1β (IL1β), cytokeratin 10 (K10), and matrix metalloproteinase-9 (MMP-9), ultimately leading to epithelial apoptosis. Persistent ROS accumulation was observed in PBS- and TCE-treated corneas, whereas a single dose of T-IDLE_STCE_ reduced ROS to near-baseline levels (Figure 7A). IL1β immunofluorescence showed intense epithelial staining in SSDE, which was profoundly attenuated by T-IDLE_STCE_ treatment; intermediate reductions were seen with TCE and eye drop therapies (Figure 7B). Elevated K10 expression, indicative of keratinization^29,30^, was markedly suppressed by T-IDLE_STCE_ (Figure 7C). MMP-9 expression, reflective of corneal barrier dysfunction^31^, was nearly eliminated by T-IDLE_STCE_, surpassing control groups (Figure 7D). TUNEL staining revealed abundant apoptotic cells in SSDE and controls, but T-IDLE_STCE_ significantly reduced TUNEL-positive cells, achieving the lowest counts among groups and reflecting enhanced epithelial repair via targeted immunomodulation (Figure 7E). Thus, T-IDLE_STCE_-mediated deep PC depletion mitigated SSDE-associated epithelial desquamation through multifaceted mechanisms, providing a superior therapeutic strategy for restoring corneal integrity and ocular surface health.

**Figure 7.**
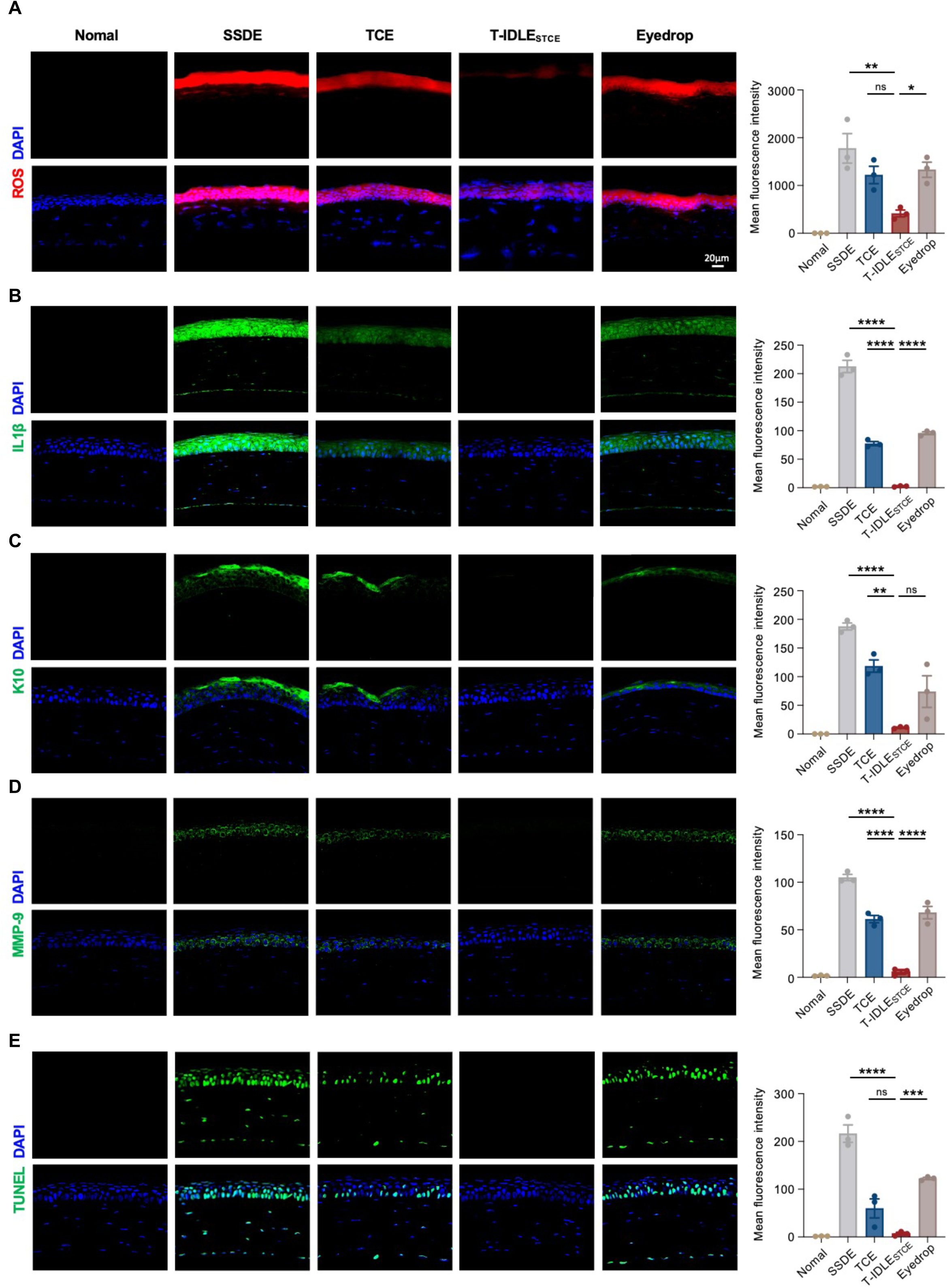
T-IDLE_STCE_ exerts anti-inflammatory, anti-apoptosis, and anti-keratinization effects in corneal tissues. (A) Immunofluorescence staining for ROS (red) in corneal tissues from each group at day 35. DAPI (blue) marks nuclei. Quantification (right) reveals ROS levels for each group (n=3 per group). Scale bar, 20 µm. (B) Immunofluorescence staining for IL1β (green) in corneal tissues from each group at day 35. DAPI (blue) marks nuclei. Quantification (right) reveals IL1β expression in each group (n=3 per group). Scale bar, 20 µm. (C) Immunofluorescence staining for K10 (green) in corneal tissues from each group at day 35. DAPI (blue) marks nuclei. Quantification (right) reveals K10 expression in each group (n=3 per group). Scale bar, 20 µm. (D) Immunofluorescence staining for MMP-9 (green) in corneal tissues from each group at day 35. DAPI (blue) marks nuclei. Quantification (right) reveals MMP-9 expression in each group (n=3 per group). Scale bar, 20 µm. (E) TUNEL staining (green) indicates apoptotic cells in corneal tissues. DAPI (blue) marks nuclei. Quantification (right) shows the extent of apoptosis in each group (n=3 per group). Scale bar, 20 µm. The numerical data in (A-E) are presented as the mean ± SEM. ns, not significant, *p < 0.05, **p < 0.01, ***p < 0.001, and ****p < 0.0001. A one-way ANOVA multiple comparisons test (A-E) was used for statistical significance analysis.

### T-IDLE_STCE_ treatment confers durable therapeutic benefits in the SSDE model

To evaluate the durability of T-IDLE_STCE_ in ameliorating SSDE symptoms, we administered a single intravenous dose of T-IDLE_STCE_ to SSDE mice at disease onset and compared with other treatments. Functional and histological outcomes were assessed at day 14 and 35 (Figure 8A). Untreated SSDE mice exhibited significantly reduced tear production, as measured by the phenol red thread test. In contrast, T-IDLE_STCE_ treatment restored tear secretion to near-normal levels by day 14 and sustained this recovery through day 35, outperforming the efficacy of both TCE and conventional eye drop treatments (Figure 8B). Corneal damage, visualized by slit-lamp fluorescein staining, was markedly attenuated in T-IDLE_STCE_-treated mice, with a significant reduction in corneal fluorescence scores compared to SSDE and TCE groups at both time points (Figure 8C). These therapeutic effects persisted through day 35, demonstrating durable epithelial protection.

**Figure 8.**
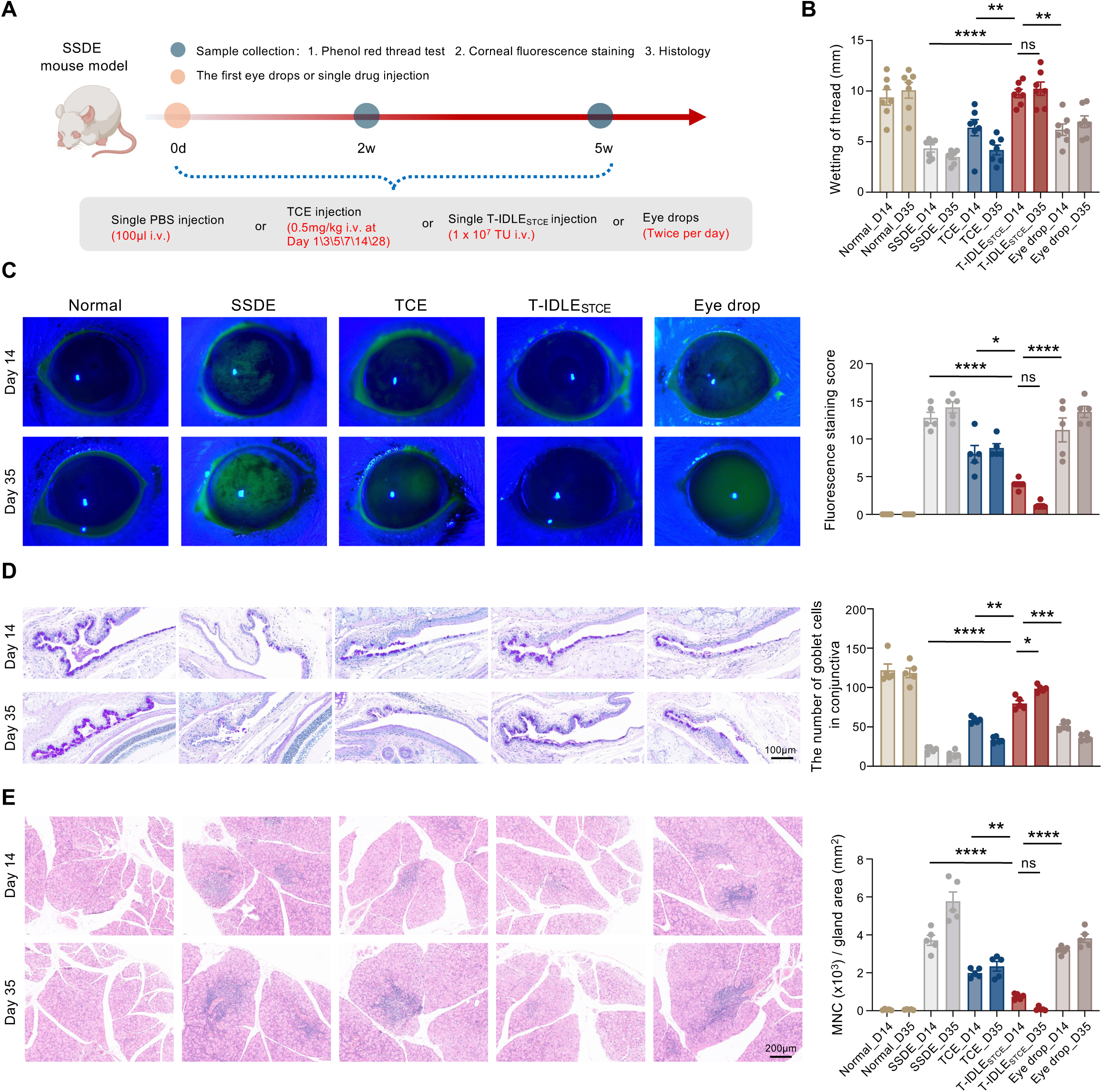
T-IDLE_STCE_ treatment exhibits long lasting therapeutic effects in SSDE model. (A) Schematic of experimental timeline. Mice received a single i.v. injection of saline, TCE (0.5 mg/kg), or T-IDLE_STCE_ (1 × 10^7^ TU), or were treated with topical eye drops (twice daily). Outcome assessments were performed at day 14 and day 35, including thread wetting test, corneal staining, and histological analysis. (B) Quantification of tear production (top) by phenol red thread test at day 14 and day 35 (n=7 per group). (C) Representative slit-lamp fluorescein images of corneas at day 14 and day 35 across groups, with quantification of staining score (n=5 per group). (D) PAS staining of conjunctival tissues at day 14 and day 35, with quantification of epithelial goblet cell numbers (n=5 per group). Scale bar, 100 µm. (E) H&E staining of ELGs at day 14 and day 35 and quantification of mononuclear-cell (MNC) infiltrates normalized to gland area (n=5 per group). Scale bar, 200 µm. The numerical data in (B-E) are presented as the mean ± SEM. ns, not significant, *p < 0.05, **p < 0.01, ***p < 0.001, and ****p < 0.0001. A one-way ANOVA multiple comparisons test (B-E) was used for statistical significance analysis.

Periodic acid-Schiff (PAS) staining of conjunctival tissue revealed significant goblet cell loss in SSDE mice; however, T-IDLE_STCE_ effectively preserved goblet cell density, whereas TCE and eye drop treatments yielded limited recovery (Figure 8D). Hematoxylin and eosin (H&E) staining of extraorbital lacrimal glands (ELGs) demonstrated dense mononuclear cell (MNC) infiltrates in SSDE glands; T-IDLE_STCE_ markedly reduced infiltration, surpassing the reductions seen with TCE and eye drops (Figure 8E).

### T-IDLE_STCE_ treatment induces potent and prolonged PC depletion in NHPs

Finally, to evaluate translational potential, we assessed the efficacy of T-IDLE_STCE_ treatment in a more clinically relevant NHP model. A single intravenous dose of T-IDLE_STCE_ (1×10⁹ TU/kg) was administered to two cynomolgus monkeys (NHP1 and NHP2), followed by monitoring of circulating PCs via flow cytometry over a period of 84 days (Figure 9A). Representative plots gated on PCs showed baseline frequencies of 19.4% in NHP1 and 28.1% in NHP2 on day 0, which declined markedly to 5.33% and 6.05% by day 14, respectively; by day 56, frequencies were 18.8% in NHP1 (partial rebound) and 4.8% in NHP2 (sustained depletion) (Figure 9B). Quantification of PC counts in peripheral blood revealed rapid depletion post-treatment, with counts dropping from baseline levels of approximately 800 cells/mL in both animals to below 200 cells/mL by day 14; NHP1 exhibited a gradual rebound starting around day 42, reaching ∼400 cells/mL by day 84, whereas NHP2 maintained low counts (<100 cells/mL) throughout the observation period (Figure 9C). Consistent with findings in murine models, T-IDLE_STCE_ treatment in NHPs elicited sustained PC depletion, supporting its translational potential for autoimmune disease therapy.

**Figure 9.**
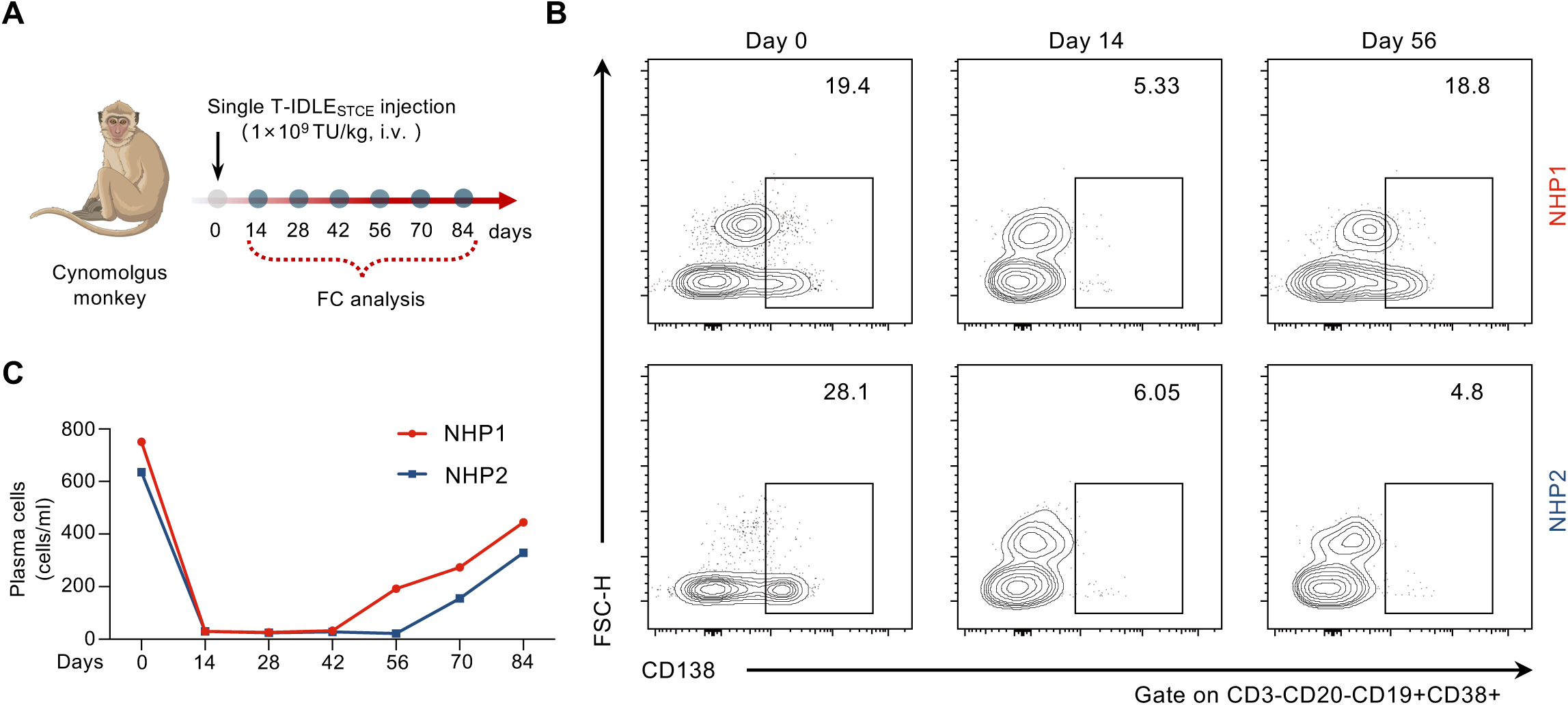
T-IDLE_STCE_ treatment induces potent and prolonged plasma cell depletion in non-human primates. (A) Schematic of the experimental timeline for *in vivo* analysis of PC depletion in cynomolgus monkeys (NHP1 and NHP2). Animals received a single intravenous injection of T-IDLE_STCE_ (1×10^9^ TU/kg). Peripheral blood samples were collected at the indicated time points (day 0, 14, 28, 42, 56, 70, and 84) for longitudinal flow cytometric analysis of circulating PCs. (B) Representative flow cytometry plots showing frequencies of PCs in peripheral blood from NHP1 (top) and NHP2 (bottom) at baseline (day 0), early post-treatment (day 14), and late post-treatment (day 56). (C) Quantification of plasma cell counts in peripheral blood over time for NHP1 and NHP2.

## Discussion

The advent of cell- and gene-based immunotherapies has transformed the landscape of cancer treatment^19,32^, and recent efforts seek to extend these advances to autoimmune diseases by targeting pathogenic immune cell populations^8,18,33^. While *ex vivo*-engineered CAR-T and TCE-T cells have demonstrated potential in eliminating autoreactive cells^13,34,35^, their widespread application in chronic autoimmune conditions remains constrained by complex manufacturing procedures, high production costs, and the risk of systemic toxicities. In this study, we introduce an *in vivo* TCE-T platform specifically designed for autoimmune disease. By engineering T-IDLE_STCE_, we convert endogenous T cells into continuous producers of a BCMA×CD3 bispecific engager. In the model of SSDE, this approach achieved superior therapeutic efficacy compared with conventional TCEs and immunosuppressive agents. The T-IDLE platform circumvents the need for *ex vivo* cell manipulation and reinfusion, mitigates infusion-related toxicities, and enables sustained drug-free remission through in-situ T cell engineering.

Compared to immunosuppressive therapies and BCMA×CD3 TCE treatments^8^, T-IDLE_STCE_ induced deeper and more durable depletion of PBs and PCs across peripheral blood, lymphoid tissues, and lacrimal glands. Importantly, STCE-T cells not only directly mediate cytotoxicity but also recruit bystander T cells to sites of pathology, promoting their expansion and preservation of effector functions. This endogenous immune activation may be crucial for sustaining disease-free states after discontinuation of treatment, which contrasts with the transient effects of traditional therapies. we propose that T-IDLE_STCE_ enhances TCE immunoregulatory potential through its distinct ability to redirect T cells and secrete bispecific antibodies, resulting in superior efficacy compared to TCE injections and CAR-T therapies^13,14^.

Another key advantage of T-IDLE_STCE_ is its ability to selectively kill pathogenic cells while the secreted TCEs shield TCR and CD3 epitopes, mitigating excessive T cell activation. In contrast to second-generation CAR-T therapies, which often induce cytokine storms due to potent co-stimulatory signals, T-IDLE_STCE_-engineered T cells release cytokines in a more controlled manner upon target engagement. This balanced activation facilitates effective depletion of pathogenic cells while avoiding off-target tissue damage and systemic immune overactivation. The primary targets of STCE-T cells are PBs and PCs, whose elimination leads to PB repopulation after 21 days, while maintaining suppressed PC levels. This approach is more precise compared to therapies targeting CD20 or CD19, with minimal impact on the body’s normal immune function. The “plasma cell reset” induced by T-IDLE_STCE_ differs from traditional cellular immune resets, yet this limited scope of immune modulation still results in a durable and effective “clinical immune reset”^8,36^. In non-human primates, T-IDLE_STCE_ induces sustained PC aplasia comparable to that reported for third-generation lentiviral vector-based *in vivo* CAR-T platforms^37^, while obviating lymph-node administration and offering a more favorable safety and operational advantages.

In summary, this study introduces a first-in-class *in vivo* TCE-T platform tailored specifically for autoimmune disease management. By converting endogenous T cells into continuous TCE-secreting cells, T-IDLE delivers deep and long-lasting clearance of pathogenic PCs, demonstrating sustained and significant efficacy. This modular, ready-to-use approach holds broad promise for diverse autoantibody-mediated diseases and has the potential to reshape cellular immunotherapy paradigms for autoimmune diseases.

### Limitations of the study

First, standardized clinical platforms for *in vivo* CAR-T therapy are still under development^38,39^, and direct comparative studies between these approaches and T-IDLE therapy are necessary to clarify relative advantages in efficacy, durability, and safety profiles. Second, while the integration-deficient design reduces the risk of insertional mutagenesis, potential off-target transduction by the vector within patients still requires long-term monitoring. Nonetheless, the compelling efficacy observed in mice and NHP provides strong justification for advancing toward phase I clinical trials, These efforts will help unlock the full clinical potential of this *in vivo* TCE platform.

## Acknowledgments and sources of funding

Ying-Feng Zheng is supported by the National Key Research and Development Program of China (2022YFC2502800), National Natural Science Foundation of China (82171034), Guangdong Basic Research Center of Excellence for Major Blinding Eye Diseases Prevention and Treatment, and State Key Laboratory of Ophthalmology, Zhongshan Ophthalmic Center at the Sun Yat-Sen University.

## Author contributions

J.L. and D.W. designed and analyzed all experiments. J.L., D.W., Jiangbo L., Jingni L., Q.H., and Q.Z. performed experiments and analyzed data. T.H. and X.L. provided advice and discussed results. J.L., D.W., Z.L., and Y.Z. wrote the paper. J.L., D.W., T.H., and Y.Z. conceptualized the project. Z.L., T.H., and Y.Z. conceived and supervised the study. All authors discussed results and commented on the work.

## Methods

### Human samples

We profiled the peripheral blood immune cell composition and soluble BCMA (sBCMA) levels in twelve patients over 18 years of age with primary Sjögren’s syndrome (pSS). Age- and sex-matched healthy donors served as controls to minimize confounding and provide a stable baseline. Blood collection and analyses were approved by the Ethics Committee of the Zhongshan Ophthalmic Center, Sun Yat-sen University (Number: 2025KYPJ095), and informed consent was obtained from all participants. All patients fulfilled the 2016 American College of Rheumatology/European League Against Rheumatism (ACR/EULAR) classification criteria for pSS. Disease activity was quantified using the EULAR Sjögren’s Syndrome Disease Activity Index (ESSDAI). Demographic and clinical information—including medical history, symptoms, and signs—was extracted from medical records. The demographic and clinical characteristics of the control and pSS cohorts are summarized in Tables S1.

### Cell lines and culture conditions

Suspension cell lines were cultured in RPMI-1640 supplemented with 2 mM L-glutamine, 10% heat-inactivated fetal bovine serum (FBS), penicillin–streptomycin (100 U/mL penicillin, 100 µg/mL streptomycin), and 0.05 mM β-mercaptoethanol (Sigma-Aldrich). Adherent cell lines were maintained in DMEM supplemented with 2 mM L-glutamine, 10% heat-inactivated FBS, and 1% penicillin–streptomycin. All cells were grown at 37 °C in a humidified incubator with 5% CO₂ and were routinely screened for mycoplasma by PCR and confirmed negative.

### Mice and NHP models

Male BALB/c and NOD/ShiLtJ mice (8 weeks old; 25-30 g) were obtained from GemPharmatech and maintained under specific pathogen-free (SPF) conditions at the Zhongshan Ophthalmic Center, Sun Yat-sen University. Animals had ad libitum access to standard chow and water. All procedures were approved by the Animal Experimental Ethics Committee of the Zhongshan Ophthalmic Center (Number: Z2025033). Two cynomolgus macaques (male, 3-3.5 years old) were singly housed in cages (height 1.0 m, depth 0.9 m, width 0.8 m) under controlled conditions (23 ± 3 °C; relative humidity 55% ± 5%). On day 0, animals received an intravenous dose of 1 × 10^9^ TU/kg. Body temperature, activity, appetite, fecal output, and overall health were monitored daily. Peripheral blood was collected on designated post-injection days for flow-cytometric analysis.

### Clinical sample collection

Peripheral venous blood (4 mL) was drawn and centrifuged to separate plasma. The remaining blood was mixed with 10 mL red blood cell lysis buffer, incubated on ice for 10 min, and centrifuged at 400 × g for 5 min. The cells were resuspended in 0.9% NaCl and carefully layered over Ficoll; peripheral blood mononuclear cells (PBMCs) were isolated by density-gradient centrifugation according to the manufacturer’s instructions. The interphase PBMCs were collected, washed, and used for downstream assays.

### Vector production

T-IDLE is a third-generation integration-deficient lentiviral vector^37^ engineered to display an anti-CD3ε single-chain variable fragment (scFv) on the viral envelope. T-IDLE_STCE_ particles were produced by transient co-transfection of HEK293T cells with four plasmids encoding gag-pol, rev, anti-CD3ε scFv-T2A-Cocal mutant envelope, and the payload vector. The sequence of the anti-mouse CD3ε scFv (clone 145-2C11) was obtained from patent US20220041724A, whereas the anti-CD3 scFv sequence in the T-IDLE envelope for use in monkeys was derived from OKT3. In T-IDLE_STCE_, the anti-CD3 scFv component of the TCE matched the aforementioned sequences, and the anti-BCMA scFv was sourced from the 25C2 clone^40^, which is cross-reactive with both mouse and human BCMA. Sixteen hours post-transfection, fresh medium containing 15 U/mL Denarase (c-LEcta) was added to the culture. The supernatant was collected the following day, clarified using Mustang Q (Cytiva), concentrated by tangential flow filtration (Sartorius Vivaflow 50, VF05P4), and passed through a 0.2 μm polyethersulfone (PES) sterile filter prior to aliquoting and storage at −80°C. An additional batch of supernatant was collected at 48 hours post-transfection, filtered through a 0.45 μm PES sterile filter, and stored similarly. A20^BCMA^ cells were generated by lentiviral transduction using BCMA-expressing vectors, in which GFP or luciferase served as reporter genes, enabling stable expression of BCMA. A20^Luc^ cells were established by transduction with a luciferase-expressing lentivirus. Both BCMA and luciferase lentiviral vectors were provided by Cyagen.

### A20^BCMA^ cell line generation

A20 cells were separately transfected with piggyBac (PB) transposon vectors EF1A-BCMA-mCherry-P2A-tdTo vector and EF1A-BCMA-P2A-luci vector including a puromycin resistance marker for further cell selection. After transduction, cells were maintained in RPMI-1640 supplemented with 10% FBS and puromycin (1 μg/mL).

### Transmission electron microscopy

Ten microliters of IDLE suspension was applied to 200-mesh carbon-coated copper TEM grids and allowed to adsorb, followed by negative staining with 2% (w/v) uranyl acetate (10 µL, 10 min, room temperature). The stain was removed, excess liquid was wicked off with filter paper, and grids were air-dried. Images were acquired on a Tecnai G2 Spirit BioTWIN (FEI) equipped with a Gatan 832 CCD camera at 150,000–250,000× magnification.

### T cell transduction

Single-cell suspensions were prepared from BALB/c mouse spleens by mechanical dissociation followed by red blood cell lysis. Primary mouse T cells were purified by negative selection using the Pan T Cell Isolation Kit II (Miltenyi Biotec) according to the manufacturer’s instructions. For transduction assay, cells were activated on plate-coated anti-CD3 (5 μg/mL; clone 145-2C11) for 4 h at 37 °C, then expanded for 2 days in medium containing soluble anti-CD28 (5 μg/mL; clone 37.51) and recombinant murine IL-2 (50 U/mL). Activated T cells were transduced with IDLEs at the multiplicities of infection (MOIs) specified for each experiment. Nontransduced primary T cells (NT-T) served as negative controls. For experiments assessing vector-mediated activation, particles were added directly to unstimulated primary mouse T cells at 1 × 10⁶ cells/mL. Activation was assessed by flow cytometry on day 1, 3, and 7. For flow cytometry, cells were first stained with a fixable viability dye, followed by surface marker staining.

To assess whether T-IDLE specifically binds T cells, DiO (10 mg mL⁻¹) was incubated with IDLE or T-IDLE at 37 °C for 10 min, after which DiO-labeled IDLEs were collected by centrifugation at 1,000 × g for 10 min. Activated primary T cells were seeded in 96-well plates (1 × 10⁶ cells per well) and incubated for 8 h with PBS, or with 1 × 10⁸ DiO-labeled IDLE or T-IDLE particles. Cells were then stained with Dil at 37 °C for 10 min and analyzed by confocal microscopy (LSM980, Zeiss). Confocal images/data were processed using ZEISS ZEN software.

### Cell-based binding assay

Single-cell suspensions of A20^BCMA^ cells and non-activated primary mouse T cells were prepared as described. Cells were labeled with CFSE or Violet (Thermo Fisher) according to the manufacturer’s instructions. Equal numbers of the two cell types were mixed and incubated on ice for 40 min with 200 μL conditioned supernatants collected on day 5 post-infection from five primary T-cell groups (PBS, IDLE, IDLE_STCE_, T-IDLE, and T-IDLE_STCE_) to assess binding mediated by T-cell secreted products. Cells were then washed twice with FACS buffer, resuspended in 300 μL FACS buffer containing DAPI, then analyzed by flow cytometry; the percentage of CFSE/Violet pairs was quantified within the live doublet gate. Single-cell suspensions of A20^BCMA-mCherry^ cells and T-IDLE_STCEGFP_ cells were prepared as described. Approximately 1×10^6^ cells of each type were seeded into 35-mm confocal dishes and co-cultured at 37 °C for 1 h. Supernatants were then removed, cells were incubated on ice for 20 min with anti-His antibody (Abcam), washed twice, and imaged by confocal microscopy. Image processing was performed using ZEISS ZEN software.

### Western blotting

Samples were resolved by SDS-PAGE under reducing conditions on 4%–20% Tris–glycine gradient gels, transferred to polyvinylidene difluoride membranes, and incubated with anti-His monoclonal antibody. To characterize IDLEs protein composition, blots were additionally probed with anti-p24 and anti-G4S linker antibodies. HRP-conjugated goat anti-rabbit IgG (H+L) and goat anti-mouse IgG (H+L) secondary antibodies were used, and signals were developed with Immobilon Western chemiluminescent HRP substrate (Millipore). Bands were imaged using the chemiluminescence imaging system (Tanon Science & Technology).

### Enzyme-linked immunosorbent assay

Human and mouse plasma soluble BCMA (sBCMA) levels were quantified using the Human BCMA/TNFRSF17 ELISA Kit and the Mouse BCMA/TNFRSF17 ELISA Kit, respectively, according to the manufacturers’ instructions. Mouse plasma IFN-γ, anti-SSA (Ro60), and anti-SSB (La) antibodies were measured by ELISA following the manufacturers’ protocols.

To detect TCE secreted into cell-culture supernatants, ELISA plates were coated overnight at 4 °C with recombinant mouse BCMA-Fc chimera (5 µg/mL). After washing and blocking, conditioned medium was added and incubated for 1 h at room temperature. Plates were washed and incubated with anti-His monoclonal antibody (1 µg/mL) for 1 h at room temperature, washed again, and then incubated with HRP-conjugated goat anti-mouse IgG (Fc-specific; 0.4 µg/mL) for 1 h at room temperature. Signals were developed with tetramethylbenzidine (TMB) substrate, and absorbance was read at 450 nm using a BioTek Synergy H1 multimode plate reader.

### Histopathology

Mice were euthanized on day 14 and 35. Whole globes and lacrimal glands were collected, fixed in 4% paraformaldehyde, and processed for hematoxylin-eosin (H&E) or periodic acid-Schiff (PAS) staining. For goblet-cell quantification, five distinct PAS-stained sections per group were analyzed, and goblet cells in the superior and inferior conjunctival fornices were counted. Corneal inflammation was evaluated by immunohistochemistry on 4% paraformaldehyde-fixed sections for TUNEL, IL1β, keratin 10 (K10), and MMP-9; reactive oxygen species (ROS) were assessed on cryosections. Lacrimal gland sections (5 µm) were subjected to immunohistochemistry for IFN-γ, IL-17, and CD3 and BCMA. For each group, ten random fields from three mice were quantified, and the area of inflammatory infiltration was measured using ImageJ. Depending on the time point, lacrimal glands from mice of the indicated ages were processed for BCMA immunohistochemistry. In addition, on day 35, major organs (heart, liver, spleen, lung, kidney, and brain) were harvested, fixed in 4% paraformaldehyde, and stained with H&E. Image analysis and quantitative measurements were performed using Image J.

### Therapeutic efficacy assessment

10-week-old NOD/ShiLtJ were randomly assigned to four groups (SSDE, TCE, T-IDLE, and eye-drop), with age- and sex-matched BALB/c mice serving as normal controls. In the TCE group, animals received tail-vein injections of TCE at 0.5 mg/kg per mouse on treatment day 1, 3, 5, 7, 14, and 28, whereas in the T-IDLE group, mice were injected via the tail vein with 1×10⁷ TU/mouse on day 1. In the eye-drop group, mice were treated twice daily with 10 µL Restasis (Allergan) per eye. On day 14, therapeutic efficacy was evaluated by clinical assessments of dry eye, including tear secretion and ocular surface parameters. Tear production was measured by the Schirmer test using phenol red cotton threads (Jingming), with threads gently placed at the middle–outer one-third of the inferior conjunctival fornix for 30 s, after which the wetted length was recorded in millimeters. Corneal epithelial integrity was assessed by fluorescein staining: 1 µL of 2% fluorescein sodium was applied to the ocular surface, followed by three artificial blinks, and corneal damage was examined under cobalt-blue illumination using a slit-lamp microscope, with staining scored by an ophthalmologist in a double-blind manner.

### Cytotoxicity assays

Activated T cells-nontransduced (NT), T-IDLE, or T-IDLE_STCE_-were cocultured, with or without freshly isolated NA-T cells, together with luciferase-expressing targets (A20^BCMA-Luc^, A20^Luc^, H929^Luc^, or K562^Luc^) at the indicated E:T ratios. After 48 h, supernatants were collected and stored at −80 °C for subsequent IFN-γ analysis. Immediately before bioluminescence readout, D-luciferin (Beyotime) was added to a final concentration of 20 µg mL⁻¹, and signal was measured on a BioTek Synergy H1 multimode plate reader. Spontaneous lysis was determined by incubating target cells with NT-T or NA-T effector cells alone. Target-cell viability (%) was calculated as the mean bioluminescence of each sample divided by that of the input number of control target cells × 100, and specific lysis was calculated relative to a baseline of 100% viability.

For noncontact Transwell cytotoxicity assays, polycarbonate inserts (0.4-µm pore size; 4.26-mm diameter; Corning) were used. Luciferase-expressing target cells (5 × 10⁴) and NA-T cells (5 × 10⁴; E:T = 1:1) were plated in the lower chamber, and A-T cells (NT, T-IDLE, or T-IDLE_STCE_) were added to the inserts at the indicated ratios. After 48 h, bioluminescence was quantified as described above.

### Flow cytometry

For in-vitro T-cell activation, primary mouse T cells transduced with IDLEs were harvested, stained with anti-CD25, anti-CD4, anti-CD8, and anti-CD69, and analyzed by flow cytometry to assess activation. To evaluate the long-term effects of IDLEs on primary T cells, cells were stained with anti-CD45, anti-CD3ε, and anti-TCRβ. For transduction efficiency in splenocytes, BALB/c mouse spleens were mechanically dissociated and subjected to red-blood-cell lysis to obtain single-cell suspensions; drugs were added at the indicated MOI to splenocyte cultures that had been activated for 2 days, and after 3 days of transduction cells were stained with anti-CD45, anti-CD3, anti-CD4, and anti-CD8a for flow-cytometric analysis. Production of STCE by T cells infected/transduced with T-IDLE_STCE_ was assessed by anti-His staining under nonpermeabilized or permeabilized conditions. Surface BCMA expression on A20, K562, and H929 cells was determined using anti-mBCMA and anti-hBCMA antibodies, respectively.

For *in vivo* transduction mice were dosed via the tail vein. At designated time points blood and draining lymph nodes and lacrimal glands were collected and dissociated then stained with anti-CD45 and anti-CD3 to assess targeting. The in-vivo clearance experiment used two panels: Panel 1-BCMA, CD3, CD45; Panel 2-MHC II, CD19, TCRβ, CD45, CD138, IgM, B220. Gating definitions were: B cells, CD45⁺ TCRβ⁻ CD19⁺; plasma cells, CD45⁺ TCRβ⁻ CD19⁻ B220⁻ CD138⁺ MHC II⁻ IgM⁺; plasmablasts, CD45⁺ TCRβ⁻ CD19⁺ B220⁺ CD138⁺ MHC II⁺ IgM⁺.

For assessing T-cell effector function in draining lymph nodes, nodes were collected at day 14 and 35 post-treatment, digested, and resuspended in medium containing Brefeldin A (1 µg/mL), PMA (50 ng/mL), and ionomycin (200 ng/mL) for 6 h, followed by staining with CD3, CD45, CD4, IL-17A, and IFN-γ. Lacrimal-gland isolation and digestion were performed as described in the literature^41^. For human peripheral-blood immune-cell phenotyping and BCMA expression, panels included CD45, CD3, CD20, CD19, CD38, CD138, and BCMA. For NHP pharmacodynamic analyses, panels included MHC II, CD19, TCRβ, CD45, CD138, IgM, and B220; B cells were defined as CD45⁺ CD3⁻ CD20⁺, and plasma cells as CD45⁺ CD3⁻ CD20⁻ CD19⁺ CD38⁺ CD138⁺. For mouse BCMA flow cytometry, dissociated single cells should be incubated in complete RPMI-1640 medium containing DAPT (10 μM) at 37 °C for 1 h, washed twice, and then subjected to surface antibody staining. All antibodies were used according to the manufacturers’ instructions, with DAPI or Zombie dyes for viability. All samples were acquired on a BD LSRFortessa (BD Biosciences, Franklin Lakes, NJ, USA) and analyzed with FlowJo software (v10.4).

### RNA seq

Lacrimal glands were harvested, immediately snap-frozen in liquid nitrogen, and submitted to BGI (BGI Genomics) for RNA-seq library preparation and sequencing. Briefly, total RNA was purified from lacrimal gland tissues and reverse-transcribed using SuperScript III reverse transcriptase according to the manufacturer’s instructions. The resulting cDNAs were fragmented by sonication on a Covaris S2 ultrasonicator, and sequencing libraries were generated with the KAPA high-throughput library preparation kit. Libraries were sequenced on the BGISEQ-500 platform. Gene expression levels were quantified as fragments per kilobase of transcript per million mapped reads. Differentially expressed genes between the indicated experimental groups were identified using DESeq2 (v1.24.0), with significance thresholds set at |log2(fold change)| > 2 and adjusted P < 0.05.

### Statistical analysis

Statistical analyses were performed using GraphPad Prism v9 (GraphPad Software). For univariate comparisons among three or more groups, one- or two-way ANOVA was applied followed by Tukey’s multiple-comparisons post hoc test. For bivariate analyses using contingency tables, statistical significance was determined by Fisher’s exact test. The specific statistical test used in each experiment is indicated in the corresponding figure legend. Significance thresholds were defined as *p < 0.05, **p < 0.01, ***p < 0.001, and ****p < 0.0001. Data are presented as mean ± SEM, and N denotes the total number of technical and biological replicates.

